# Characterization of the mesendoderm progenitors in the gastrulating mouse embryo

**DOI:** 10.1101/2024.04.28.591221

**Authors:** V. Pragathi Masamsetti, Nazmus Salehin, Hani Jieun Kim, Nicole Santucci, Megan Weatherstone, Hilary Knowles, Jane Sun, Riley McMahon, Josh B. Studdert, Nader Aryamanesh, Ran Wang, Naihe Jing, Pengyi Yang, Pierre Osteil, Patrick P.L Tam

## Abstract

A population of putative mesendoderm progenitor cells that can contribute cellular descendants to both mesoderm and endoderm lineages is identified. These progenitor cells are localized to the anterior primitive streak and the adjacent epiblast of E7.0-E7.5 mid-to late-gastrula stage embryos. Lineage tracing in vivo revealed that putative mesendoderm progenitors that are marked by *Mixl1* and *Mesp1* activity contribute descendants to the endoderm layer. Analysis of the role of Mixl1 transcription factor in endoderm differentiation of the mouse epiblast stem cells revealed the choice for endoderm or mesoderm cell fate depends on the timing of activation of *Mixl1* upon exit from pluripotency, suggesting Mixl1 function may underpin the divergence of the mesendoderm progenitor to mesoderm and endoderm lineages. The knowledge gained on the spatial, temporal, and lineage attribute of mesendoderm progenitors enriches our mechanistic understanding of germ layer allocation and endoderm differentiation of mesendoderm progenitor in embryonic development and lineage allocation of primed state pluripotent stem cells in vitro.

## Introduction

During mouse gastrulation, the formation of germ layers proceeds concurrently with the transition of pluripotency state and the allocation of the multipotent epiblast cells to the germ layer derivatives [1–4]. Lineage analysis and fate-mapping studies have revealed that groups of cells in different regions of the epiblast and the primitive streak of the gastrulating embryo are allocated to progenitors of the three germ layers with descendants of each progenitor contributing to different types of germ layer derivatives [5–11]. Spatiotemporal transcriptomic analysis of the epiblast cell population has inferred that, besides the allocation of the epiblast cells to separate progenitors of ectoderm, mesoderm and endoderm, a population of putative bipotent mesendoderm progenitors may be present in the gastrulating embryos. Consistent with the functional ontology of genes enriched in these progenitors, the descendants of these progenitors are predicted to contribute to the anterior mesoderm, nascent mesoderm, definitive endoderm and axial mesendoderm [12].

Whether these bipotent mesendoderm progenitors are present in the gastrulating embryo has not been established. Using a dual Brachyury(T)/Sox17 reporter, it has been previously shown that a population of *T*^low^/*Sox17*+ cells was detected among the cells in the epiblast and the primitive streak of the mouse embryo and the stem cell-based gastruloids, suggesting that some cells in the epiblast and primitive streak may have dual germ layer potential and function as the mesendoderm progenitor [13]. Genetic tracing studies of the specification of anterior mesoderm and definitive endoderm at gastrulation showed that these two lineages emerge from a population of *Eomes*-expressing progenitors early in gastrulation, followed by the segregation of the *Eomes*-expressing cell population into *Eomes+/Mesp1+* mesoderm progenitors and *Eomes+/Foxa2+* endoderm progenitor[14]. Analysis of co-expression of lineage markers for endoderm (FOXA2) and mesoderm (Mesp1venus) in the *Eomes*-expressing cell population revealed that triple marker *(Eomes/Mesp1/Foxa2*)-positive cells, the putative mesendoderm progenitors, are present in the late-gastrulation embryo, and they may represent an intermediate cell type that emerges transiently during lineage segregation [14].

Putative mesendoderm progenitor have been identified in pluripotent stem cells during lineage differentiation in vitro. By inducing mouse embryonic stem cells (ESCs) with Activin or Nodal, cells expressing *Gsc*, E-cadherin and PDGFRa could be produced in vitro. These cells behave like mesendoderm progenitors in that single cells from which can differentiate into both endoderm and mesoderm cells in vitro [15]. In the human ESCs, a transient population of four factors *EOMES/MESP1/MIXL1/T*-expressing cells were detected at Day 2 of differentiation to the cardiomyocyte lineage [16]. Mouse epiblast stem cells (EpiSCs) derived from the epiblast of gastrulating embryos display enhanced lineage propensity for mesoderm and endoderm, reminiscent of that of mesendoderm progenitors and typically express *T* and *Mixl1* at the early phase of lineage differentiation in vitro [17]. The co-expression of *T* and *Mixl1* is reminiscent of the co-expression of Xbra and Mixl1 in the marginal zone of the *Xenopus* embryo where progenitors of mesoderm and endoderm are co-localised [18]. While the role of Mixl1/MIXL1 in driving endoderm differentiation in stem cells is yet unclear, that these genes are expressed during mesendoderm differentiation suggests *Mixl1/MIXL1* may be employed as an additional marker for mesendoderm progenitors. In this study, we sought to characterize the spatiotemporal localization of the putative mesendoderm progenitors (pMEP) in the gastrulating mouse embryo by capitalizing on the knowledge of the association of *Mixl1/MIXL1* with mesendoderm differentiation of the human ESCs and mouse EpiSCs.

In the mouse embryo, *Mixl1* is first expressed in the primitive streak cells and the neighboring epiblast cells, but not in the definitive endoderm [19]. MIXL1/Mixl1 activity has been associated with the differentiation of the human and mouse pluripotent stem cells into a transient population of primitive-streak like cells, which co-express *MIXL1/Mixl1* and *OCT4/Oct4*, that comprise mesoderm and endoderm precursors [20]. *Mixl1* is also expressed in the putative MEPs induced from the mouse ESCs [15] and the primitive-streak like cells derived from hESCs that constitute the primitive haematopoietic precursors [21]. Enforced expression of *Mixl1* in mESCs, however, suppresses haematopoietic mesoderm differentiation while promotes endoderm differentiation [22]. Loss of *Mixl1* activity leads to deficiency of anterior definitive endoderm and axial mesendoderm but does not impact on mesoderm formation [23] . Since *Mixl1* activity is absent in the definitive endoderm, this phenotypic outcome may suggest that *Mixl1* may play a role in mediating the differentiation of the endoderm progenitors while they are in the primitive streak. Consistent with the notion that *Mixl1* activity is associated with mesendoderm precursors, the expression of Mix.1, the homologue of *Mixl1*, is induced in the presumptive endoderm in the animal cap tissue of the Zebrafish embryo [24] and in the precursors of anterior endomesoderm of the *Xenopus* embryo [25]. In this study, we investigated the functional attributes of Mixl1 in guiding the differentiation of the putative MEPs in the EpiSC model, in order to gain mechanistic insights into the lineage segregation of the mesendoderm progenitors.

## Results

### Identifying the mesendoderm cells in the gastrulating embryo

To infer the location of putative mesendoderm progenitors within the gastrulating embryo, we computed the activity scores of the mesendoderm genes: *T/ Brachyury* and *Mixl1*, and lineage-related genes: *Mesp1, Rbm47, Foxa2 and Sox17*. The activity score mapped the location of cells co-expressing these genes by reference to the spatiotemporal transcriptome (E5.5, E6.0, E6.5, E7.0 and E7.5 embryos [26], to the primitive streak and adjacent epiblast of E7.0 embryos (Figure S1A). To identify cells with dual lineage potential, we performed single cell RNA seq on whole E7.0 and E7.25 embryos and posterior halves of E7.5 embryo (Figure S1B). We recovered a total of 5886, 2683, 4869 cells from E7.0, E7.25 and E7.5 embryos respectively (Figure S1C) from which we constructed a reduced dimension representation of the data, using a uniform manifold approximation and projection (UMAP) (Figure 1A), and annotated inferred cell types based on differentially stably expressed genes (Figure S1D) [27]. The results were visualized on UMAP representations of cell types by time stamps of the samples (Figure 1A). The cell types identified in our dataset were compared to a published dataset of the gastrulating E7.5 mouse embryo (GSE121708) (Figure S1E). Similar cell types from both datasets clustered together and showed high correlation scores. We identified additional cell types in our dataset, e.g., primitive streak, cardiac progenitors, epiblast, EXE ectoderm and visceral endoderm, likely due to the sampling of earlier time points (E7.0 and 7.25) in our dataset other than a single time point at E7.5 (GSE121708).

**Figure 1:**
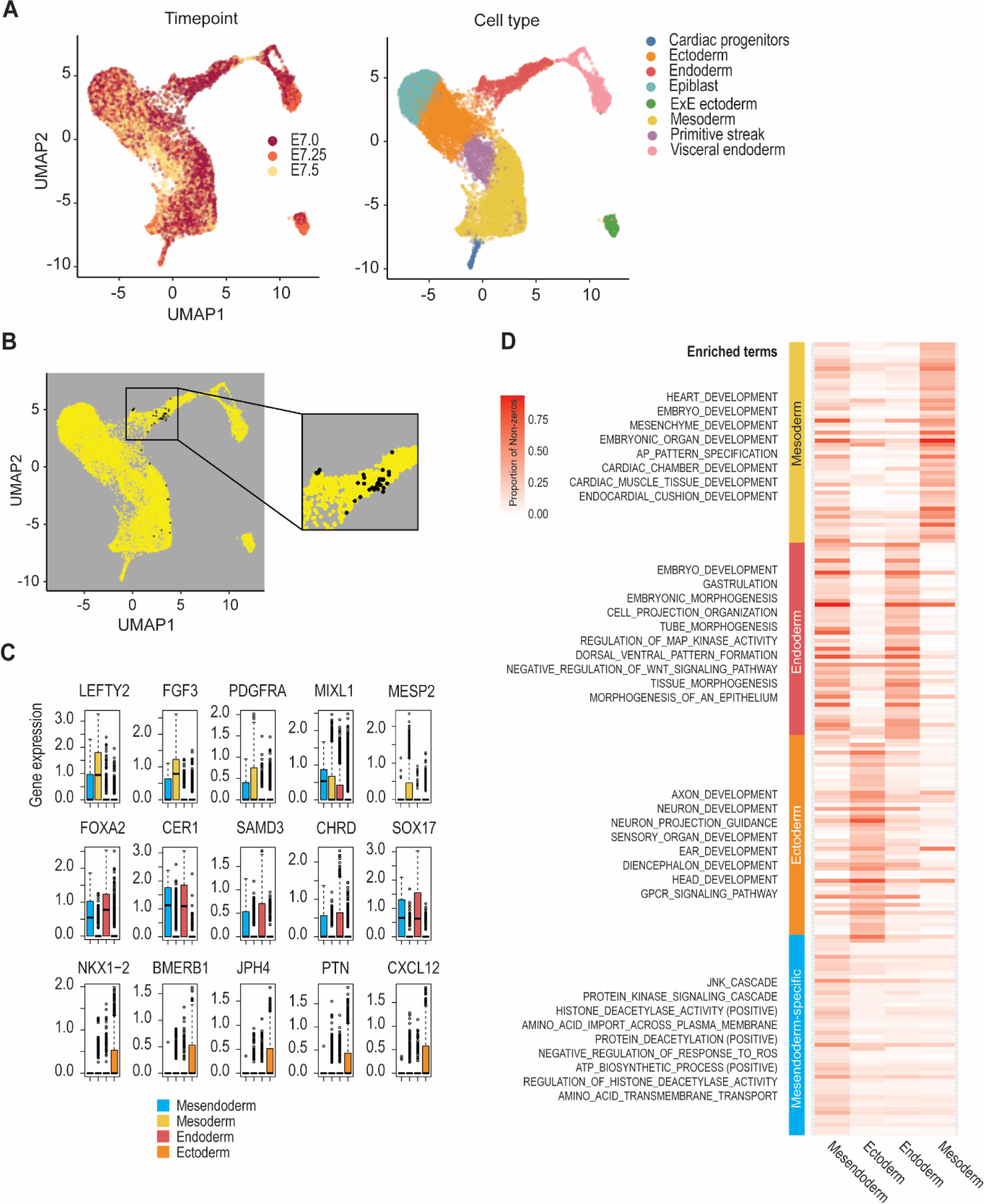
**Identification of Mesendoderm progenitor population in mouse embryos.** A. UMAP representation of single cells sampled from the gastrulating embryos coded by developmental time points (left) and cell types (right). B. UMAP representation of putative mesendoderm progenitor cells (black). C. Boxplot of expression profiles of top markers of the mesoderm (top row), endoderm (middle row), and ectoderm (bottom row) lineages in the mesendoderm, mesoderm, endoderm, and ectoderm cells. D. Heatmap of the proportion of cells expressing the top 50 cell type-related markers and the enriched gene ontology term in mesendoderm, ectoderm, mesoderm and endoderm

We reasoned that cells with stable expression of both mesoderm and endoderm marker genes are putative dual potential progenitor cells (Figure 1B). In total, 44 cells co-expressing mesoderm and endoderm-related genes (Figure S1F), were identified in the clusters of mesoderm, primitive streak, and endoderm cell types and in the posterior distal epiblast of the embryo (Figure 1B). Gene expression profiles of top markers of the mesoderm, endoderm, and ectoderm lineages were examined in four cell populations, mesendoderm, mesoderm, endoderm, and ectoderm cells. (Figures 1C, D, S2A, B). While *Lefty, Fgf3* and *Pdgfra* were expressed in the mesoderm and *Foxa2, Cer1, Samd3, Chrd* and *Sox17* were expressed in the endoderm, the mesendoderm expressed all eight of the genes. We further analyzed and visualized the proportion of cells expressing the top 50 mesoderm, endoderm, ectoderm, and mesendoderm-related markers (rows) in the four cell types (Figure 1D). Confirming our analysis, the top 50 marker genes of mesoderm and endoderm were highly expressed in the respective cell type clusters, with the mesendoderm genes expressed in both the endoderm and mesoderm cell clusters.

The identification of putative mesendoderm cells allowed us to further define a set of genes that are uniquely upregulated in the mesendoderm (Figure 1D). We identified in the single cell dataset of the primitive streak (728) and mesoderm cells (1315) of posterior halves of the E7.5 embryo, a set of top mesendoderm specific genes which, in addition to *Mixl1* included *Bdh1, Enpp3, P1h1d2, Tmem231* and *1700094d03rik* (Figure 1D). The combined activity score of this gene set enabled the inference of the location of the putative mesendoderm progenitors to the primitive streak and adjacent epiblast of the E7.5 embryo of the gastrulating stages (Figure 2A), consistent with our previously inferred location of mesendoderm cells (Figure 1A). We further defined a set of genes that are highly correlated with *Brachyury/T* expression, a primitive streak regulated marker gene at E7.0 using the iTranscriptome database [26], as the spatial allocation of mesendoderm gene set pointed to primitive streak region. Comparing the spatial pattern of genes that are co-expressed with *Brachyury/T* in gastrulating mouse embryo has pinpointed pattern 4 of E7.0 embryo being the most correlated gene expression pattern (Figure S3A, B, Supp. Table 1). We then compared the mesendoderm specific gene set identified in this study and GSE121708 to genes expressed in pattern 4 of the E7.0 embryo and identified a common set of 20 genes (Figure 2B, C). This enriched mesendoderm gene set primarily expressed in the primitive streak, where we predict the bulk of the putative mesendodrm progenitors (pMEP) may be localized in E7.0, E7.25 and E 7.5 embryos (Figures 2D, S3C). Using the data from MouseGastrulationData [28], we predicted that the *Mixl1*-expressing cells could contribute to the endoderm cells in the embryonic gut (Figure 2E). The findings further affirmed that *Mixl1* may be a robust marker for the mesendoderm progenitor.

**Figure 2:**
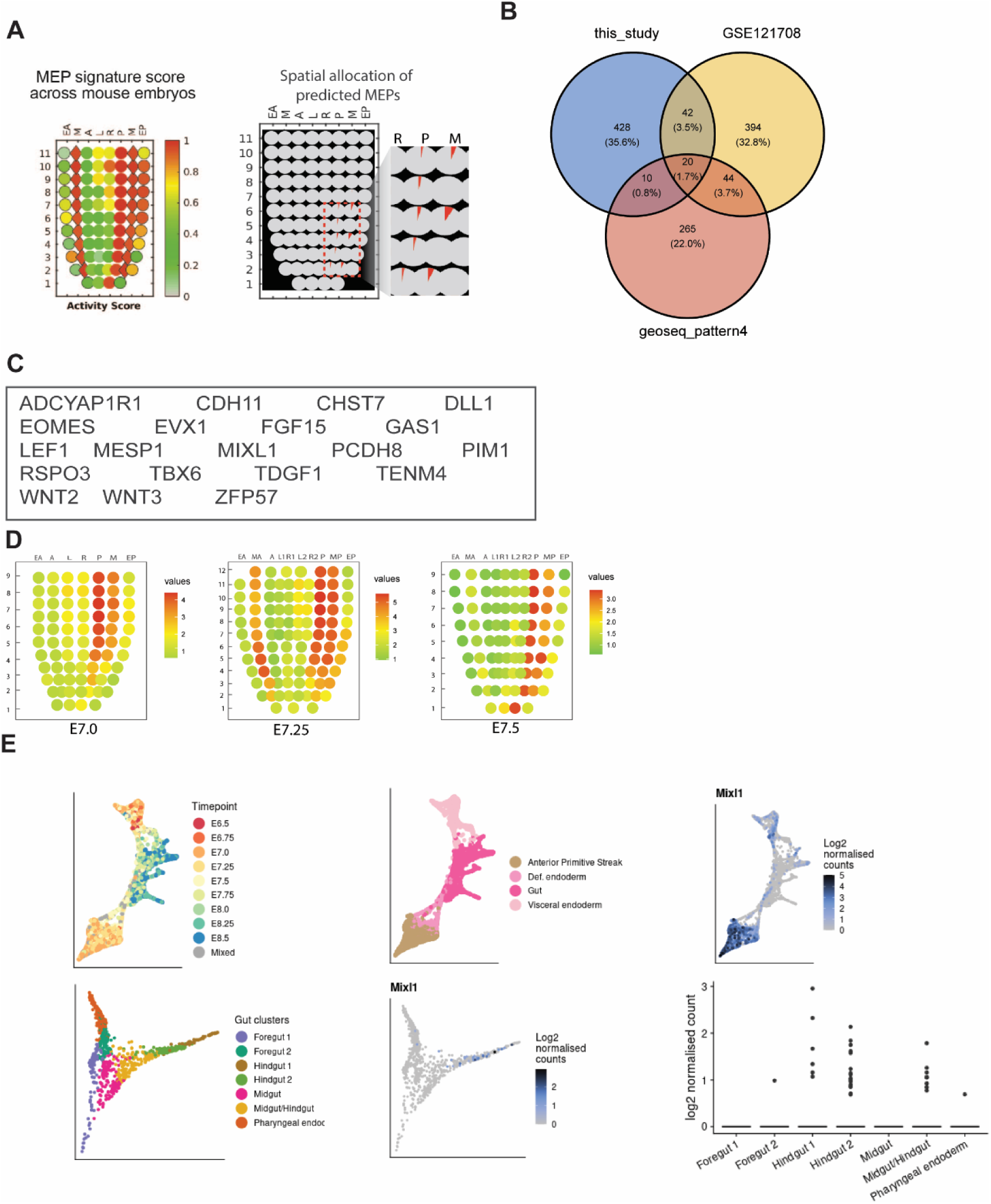
**Characterization of putative mesendoderm progenitor cells** A. Activity score of mesendoderm signature genes and mapping the spatial location of the putative mesendoderm progenitors (red sector as proportion of total cells) in E7.5 mouse embryo. B. Overlapping sets of genes between this study, the public dataset GSE121708 and Geo-seq spatial transcriptome. C. The mesendoderm related genes represent the common genes of the three datasets in (B). D. The location of the putative mesendoderm progenitor, inferred by the activity scores of the common mesendoderm-related genes in the germ layers of E7.0, 7.25 and E7.5. E. Lineage contribution of *Mixl1*-expressing cells in E6.5 to E8.5 embryos (top row) and in the gut tissue clusters of E8.5 embryo (bottom row), inferred from single-cell transcriptome dataset (Gastrulation Atlas, Pijuan-Sala et al. 2019).

### Lineage tracing of putative mesendoderm progenitor

Having identified *Mixl1* as a mesendoderm progenitor marker using single cell transcriptomics of mouse embryos, we sought to identify the descendants of *Mixl1* expressing cells *in vivo. Mixl1* is expressed in the primitive streak cells of E7.0 to E7.5 embryo and the mesoderm of E7.5 embryo but is downregulated in the endoderm layer of the embryo [19]. To track cells that had expressed *Mixl1* in the short-term, we took advantage of the Delmix (*Mixl1^+/GFP^*) mouse line, in which the *Mixl1* locus harbors a knock-in GFP reporter replacing the *Mixl1* coding sequence [23]. During gastrulation and immediately after, GFP persists in the endoderm due to the perdurance of the GFP protein (Figures 3A, B), thus allowing the use of this reporter in tracing the descendants of the *Mixl1*-expressinng pMEPs. From the Delmix x ARC cross, Mixl1-GFP expressing embryos at E7.0 to E7.5 were harvested for imaging analysis to visualize the location of Mix1-GFP positive cells in the embryo in optical Z-stacks at single cell resolution (Figures 3A, S4A, B, Supp. Movie 1). GFP positive cells could be found in the primitive streak, mesoderm and endoderm of E7.5 embryos (Figure 3B, Figure S4B). A subset of these in the endoderm and adjacent anterior primitive streak, mesoderm and endoderm. Some Mixl1-GFP positive cells in the endoderm co-expressed FOXA2, like the cells in the adjacent anterior primitive streak (Figures 3C, S4C, Supp. Movie 2). These cells were distinguishable from the resident Sox17-expressing visceral endoderm cells (Figure S4C). MIXL1-GFP positive cells were found to distribute congruently from the posterior to the anterior primitive streak as the embryo developed from E7.25 to E7.75 and contributed to the mesoderm in E7.75 embryos (Figure 3D). By E7.75, GFP/FOXA2 positive cells were detected in the definitive endoderm population (Figures 3E, S4D, E). In contrast, MIXL1GFP positive cells in the mesoderm layer did not express either FOXA2 or SOX17 endoderm marker (Figure S4D).

**Figure 3:**
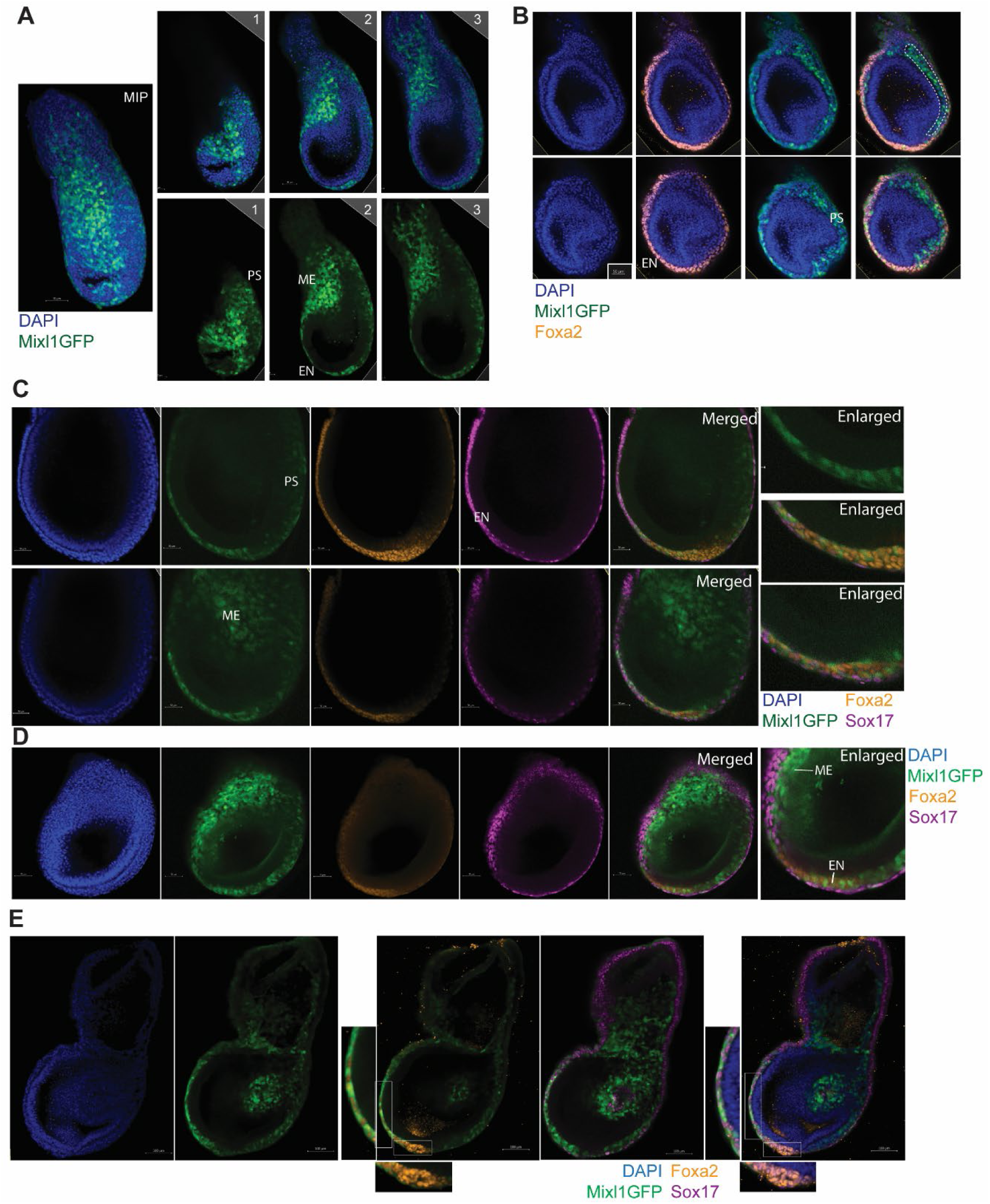
**Localization of putative mesendoderm progenitors and descendants in the gastrulating embryos** A. Mixl1GFP expressing cells in whole E7.25 embryo (at maximum intensity projection, left panel) and in the primitive streak, nascent mesoderm and endoderm (z-slices images right panel). DAPI (blue) and Mixl1GFP (green). B. Representative z-slice images of the E7.5 embryo showing Mixl1GFP expressing cells in the primitive streak (PS) and among the FOXA2+ cells (orange) in the anterior endoderm (EN). Scale bar, 50µm. C. Representative images of mouse embryo showing distribution of Mixl1GFP cells in the E7.5 embryo: the endoderm layer (EN), primitive streak (PS), and the mesoderm (ME). Top panel and bottom panel are two z-slices of the same embryo. Enlarged sections represent the cells with co-expression of Mixl1 and Foxa2 in the endoderm layer. Scale bar, 50µm. D. An example of mouse embryo in transverse z-slices showing distribution of Mixl1GFP+ cells in the mesoderm (ME) and the anterior endoderm (EN) of the E7.5 embryo. Mixl1GFP (green), Foxa2 (orange), Sox17 (purple) and DAPI (Blue). C and D are representative images from N=10 embryos. E. Representative image of E7.75 mouse embryo showing distribution of Mixl1GFP+ cells and presence of Mixl1GFP+/Foxa2+ cells in the anterior midline mesendoderm (enlarged images) but not into Foxa2 (orange) in the node region (N= 3). Scale bar, 100µm.

*Eomes*-expressing cells in the primitive streak are reputed to be the precursors for cells expressing *Mixl1* and *Mesp1* [29]. In addition to *Mixl1*, *Mesp1* is also a marker gene of the pMEPs (Figure 1C). Lineage tracing analysis performed on MouseGastrulationData [28] predicted that *Mesp1*-expressing cells in the anterior primitive streak may contribute to the endoderm in different regions of the embryonic gut (Figure S5A). To trace the lineage contribution of the *Mesp1*-expressing cells in vivo, we employed Mesp1Cre::ZEG reporter mice from which GFP-expressing embryos were harvested at E7.0, E7.5, E8.5 and E9.5 [30] (Figure S5B). As previously reported, cells that are persistently marked by Mesp1Cre-activated GFP-reporter were detected in the mesoderm layer and the heart (Figure S5B) [31–33]. A small population of GFP positive cells co-expressing SOX17 were present in the endoderm layer of E7.5 embryo (Figure 4A). By E8.5, SOX17-positive GFP-expressing cells (Figure 4B), as well as FOXA2-positive GFP-expressing cells were found in the foregut endoderm (Figures 4C, S5C, D, Supp. Movie 3) and the open mid-gut region of the embryo (Figure 4D). These findings suggest that *Mesp1*- and *Mixl1*-active progenitor cells contribute to both mesoderm and endoderm in the developing embryo.

**Figure 4:**
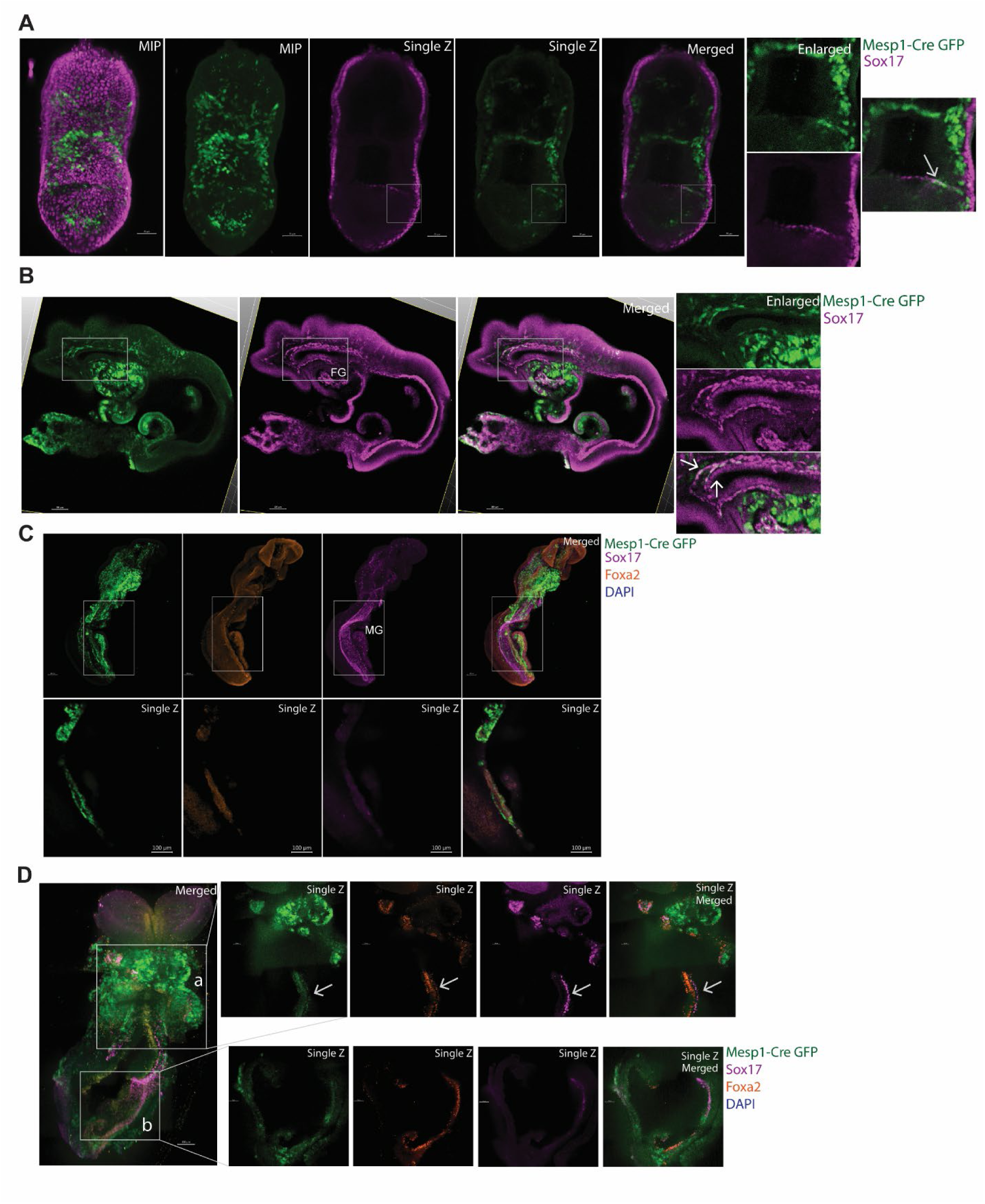
**Location of the descendants of putative mesendoderm progenitors in late and post-gastrulation embryos** A. Representative images with presence of GFP+/Sox17+ cells in the endoderm of the E7.5 Mesp1Cre::GFP embryo. GFP represents the lineage of Mesp1 expression. Enlarged images show a few cells in the endoderm (arrows) co-expressing GFP and Sox17 (purple). N=3 samples. B. Representative images with presence of GFP+/Sox17+ cells in the foregut of E8.5 Mesp1Cre::GFP. GFP represents the lineage of Mesp1 expression. Enlarged images (single z-slice) show cells co-expressing (indicated by arrows) GFP and Sox17 (purple) in the rostral-dorsal part of the foregut (FG). N=3 samples. C. Presence of GFP+/Sox17+/Foxa2+ cells in the gut endoderm of the E8.5 Mesp1Cre::GFP mouse embryo (MIP). Enlarged images (in the lower panel) show (single z-slice) the co-expression (yellow) of GFP (green), FOXA2 (orange) and SOX17 (purple) in the endoderm in the open part of the midgut (MG). Scale bar, 100µm. D. Representative image showing the presence of GFP+/Sox17+/Foxa2+ cells in the gut endoderm of the more advanced E8.5 Mesp1Cre::GFP mouse embryo. Enlarged images (single z-slice) show the co-expression (yellow), indicated by the arrow, of GFP (green)-, Foxa2 (orange)- and Sox17(purple)-positive endoderm cells in two regions (a, b) of the open midgut. Scale bar,100µm.

### Identifying the presumptive mesendoderm progenitor by MiXL1/Mixl1 expression in human embryonic stem cells and mouse epiblast stem cells

We sought to identify cells expressing multiple lineage specific markers during *in vitro* differentiation of stem cells towards primitive streak cell analogues. To drive human ESC towards the primitive streak-like cells, HES3 human ESCs harboring a transcriptional MIXL1 reporter (HES3 MIXL1-GFP) [34] were cultured as 2D micropatterns and treated either with BMP4, CHIR or CHIR and Activin (Figure 5A). Cells started expressing MIXL1GFP 24 hours after BMP4 treatment, 42 hours after CHIR treatment and 30 hours after CHIR+Activin treatment (Figures 5B, S6A, Supp. Movies 4-6). Compared to CHIR+Activin treatment, BMP4 or CHIR treatment induced fewer Mixl1GFP cells (Figures 5B-D, S6B-E). After 48 hours in CHIR+Activin, two subtypes of MIXL1GFP expressing cells were detected: one that co-expressed MIXL1 and Brachyury/TBXT, which may be the mesoderm derivatives (Figure S6D) and the other which co-expressed MIXL1 and FOXA2, representing endoderm derivatives (Figure 5D). Further analysis of the FOXA2 co-expressing cells within the CHIR+Activin condition revealed the presence of FOXA2+/SOX17+ definitive endoderm cells among the MIXL1-GFP+ cells, confirming the emergence of definitive endoderm (Figures 5D, E, S6F-H). In comparison to CHIR +Activin, CHIR treatment alone induced fewer MIXL1GFP expressing cells and MIXL1 and FOXA2 co-expressing cells, suggesting that a combination of Wnt and TGFβ signaling is required to induce robust MIXL1 expression and endoderm differentiation. In contrast, hESCs treated with BMP4 showed a preponderance of MIXL1GFP+/T+ cells, but very few MIXL1GFP+/FOXA2+ or MIXL1GFP+/SOX17+ cells, (Figures 5E, S6H), suggesting that BMP treatment alone does not is insufficient for endoderm differentiation. However, all three conditions confirmed, using an invitro model of gastrulation, that mesoderm and endoderm cell types descend from the MIXL1-expressing population.

**Figure 5:**
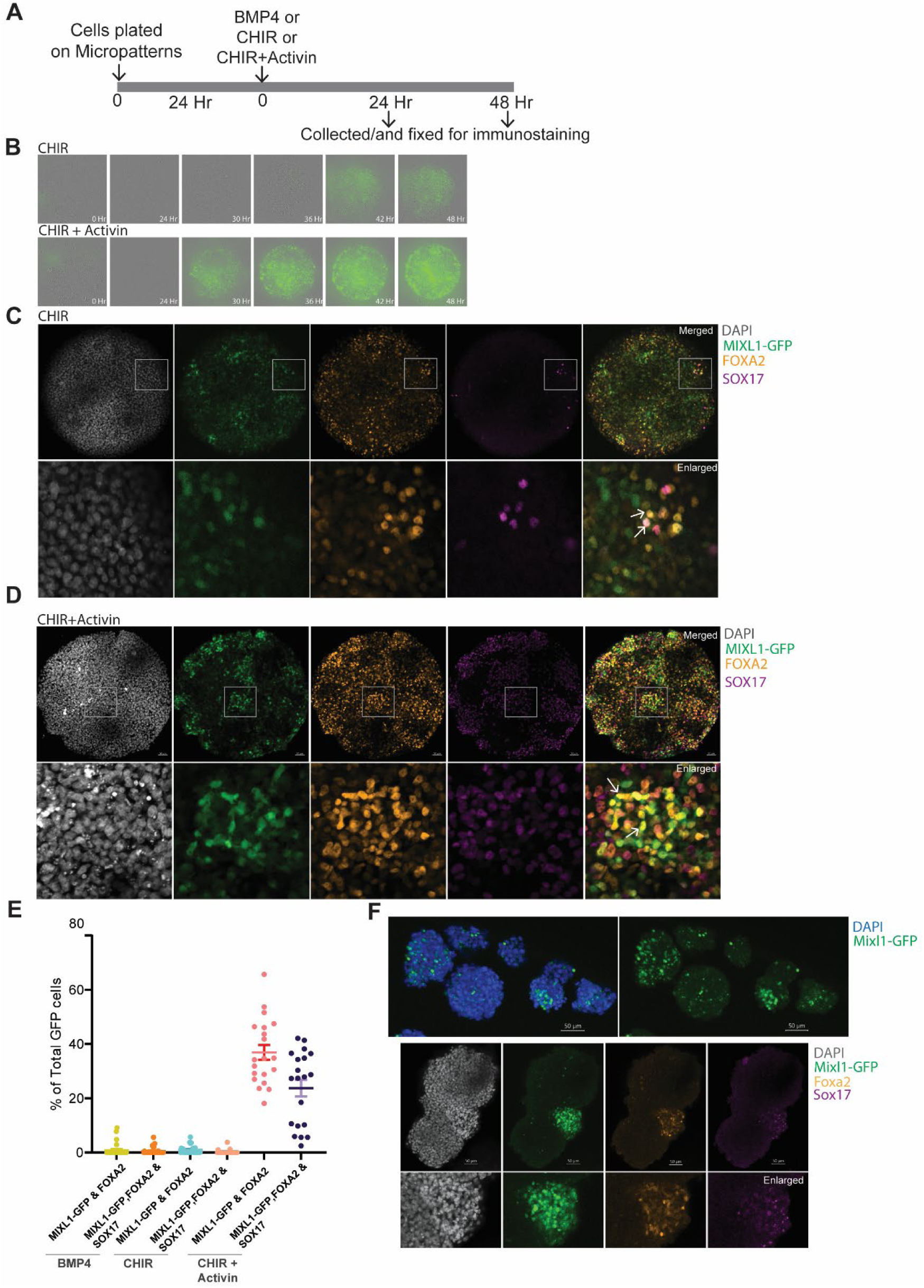
**Mesendoderm-like cells in stem cell-derived micropatterns and embryoid bodies.** A. Timeline schematic of HES3 MIXL1: GFP human ESCs differentiated into 2D micropatterns. BMP4 or CHIR or CHIR + Activin were added to the media 24 hours after plating cells. Micropatterns are collected for immunostaining after 48 hours of treatment with BMP4, CHIR or CHIR +Activin. B. Live images visualized using IncuCyte of HES3 MIXL1: GFP human ESCs in micropatterns at 0 to 48hr after treatment of CHIR alone or CHIR +Activin. C. Representative image of micropatterns (N= 30 in three independent replicates) treated with CHIR for 48 hrs. Enlarged section represents single cell resolution of co-expression (yellow) of MIXL1GFP (green), FOXA2 positive (orange) and SOX17 positive (purple) cells. Arrows point to MIXL1-GFP/FOXA2 /SOX17 positive cells. D. Representative image of micropatterns (N= 30 in three independent replicates) treated with CHIR + Activin for 48 hrs. Enlarged section represents single cell resolution of co-expression (yellow) of MIXL1GFP (green), FOXA2 positive (orange) and SOX17 positive (purple) cells marking the definitive endoderm. Arrows point to MIXL1-GFP/FOXA2/SOX17 positive cells (yellow). Scale bar, 50 µm. E. Quantification of cells co-expressing MIXL1 with FOXA2 and/or SOX17 in micropatterns following BMP4, CHIR and CHIR + Activin treatment. F. Embryoid bodies generated from Mixl1-GFP mouse epiblast stem cells showing expression of Mixl1-GFP (green), Foxa2 (orange) and Sox17 (purple). The lower panel of images show the cells with co-expression of Mixl1, Foxa2 and Sox17. DAPI in grey. Scale bar, 50 µm.

We next examined the primed mouse stem cells that are analogous to human ESCs if endoderm is derived with the *Mixl1*-expressing cell population. Our previous work on mouse epiblast stem cells (EpiSCs) showed that expression of *Mixl1* at the initial phase of in vitro differentiation enhanced the propensity for endoderm differentiation [17]. To track the emergence of *Mixl1* expressing progenitor cells, EpiSCs harboring the Mixl1-GFP reporter were generated by *in vitro* conversion of the Mixl1-GFP mouse ESCs (Figure S7A-C). Upon differentiation of the EpiSCs to embryoid bodies, Mixl1GFP+ cells emerged (Figure 5F), followed later by the appearance of Mixl1GFP+/FOXA2+ cells, MixL1-GFP+/FOXA2+/SOX17+ endoderm cells and FOXA2+ mesendoderm-like cells (Figure S7D, E). The Mix1GFP cells may therefore be an intermediate cell type in the trajectory of mesendoderm differentiation of EpiSCs.

### Timing of *Mixl1* activation predisposes the propensity of endoderm differentiation and guides cell fate choice

We next tested the functional attribute of *Mixl1* activity in eliciting endoderm differentiation in mouse EpiSCs, which display the propensity of mesendoderm differentiation reminiscent of the epiblast and primitive streak cells of the mid-(E7.0) to late primitive streak stage (E7.5) embryo [17, 35]. To generate loss-of-Mixl1 function EpiSCs, two unique biallelic frameshift mutations were introduced in the first exon of Mixl1 in A2Lox.Cre ESCs [36] (Figure 6A). Codon mapping of the transcripts predicted that these mutations produced both a premature STOP codon and inserts in the homeobox region resulting in Mixl1 loss-of-function (LOF). The LOF ESCs and the parental cell line (Figure 6A) were used to derive the EpiSC lines via the chimera detour approach [37]. Upon serum induction of differentiation, EpiSCs with intact *Mixl1* (Mixl1WT) activated *Mixl1* expression which peaked at Day 2-3 of culture, whereas the *Mixl1* knock-out EpiSCs (Mixl1KO) showed no activation (Figure 6B, S8A).

**Figure 6:**
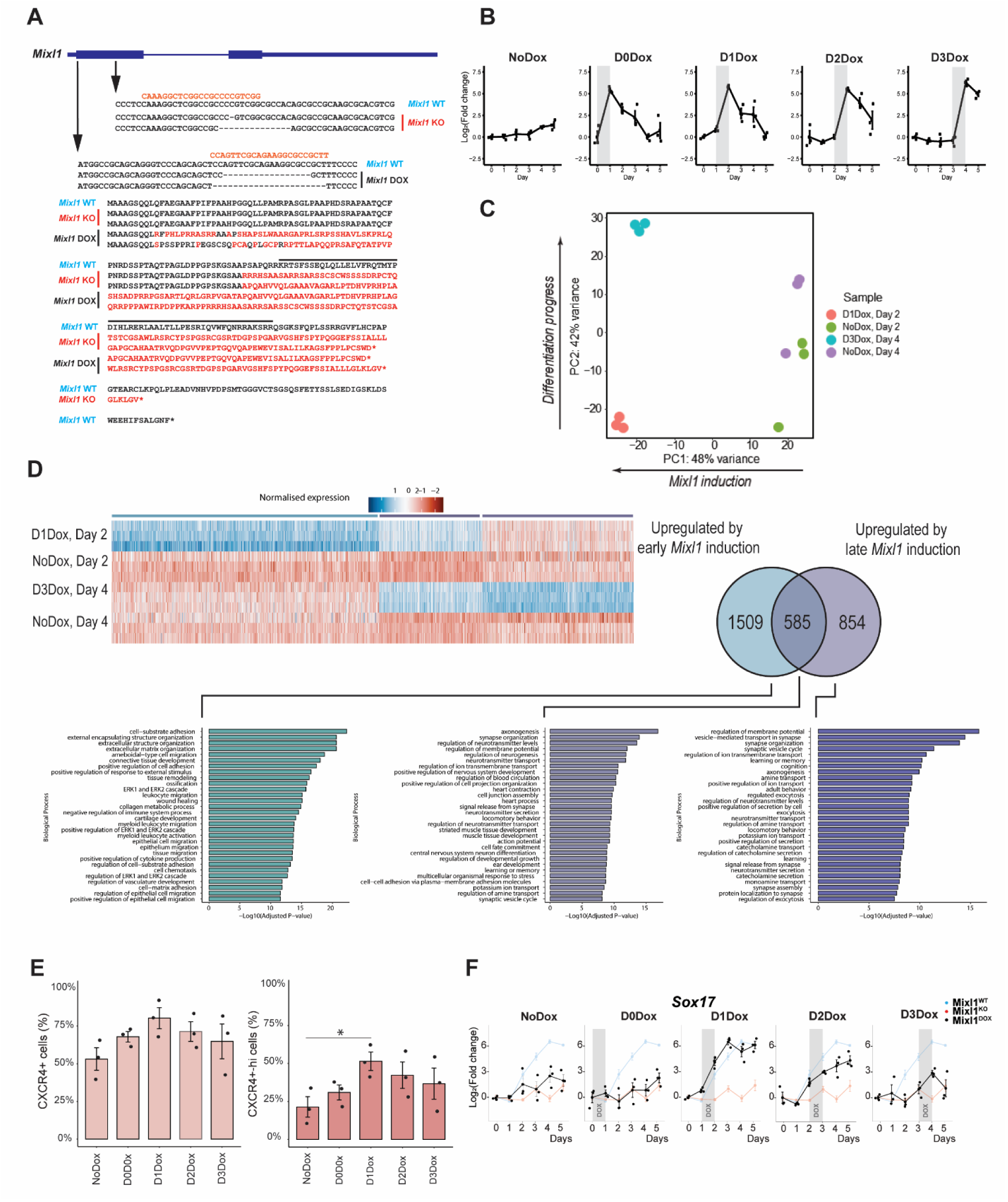
Timing of Mixl1 expression influences the choice of cell fates. A. EpiSC lines of Mixl1WT and Mixl1KO genotypes, and harboring FLAG-Mixl1 in HPRT locus that can be activated by induction with Doxycycline. Gene editing of Mixl1 locus in Mixl1 KO and Mixl1 Dox inducible cell line. B. The expression of Mixl1 in the cell lines induced by Doxycycline treatment on Day 0, 1, 2 and 3 (D0DOX, D1DOX, D2 DOX and D3 DOX respectively).The panel highlights (grey) the time when expression peaks for individual Dox induction timings. C. Principal component analysis of gene expression profile assayed by RNA sequencing of the samples-D1Dox (Day 2), NoDox(Day 2), D0Dox (Day4), NoDox (Day 4). D. Heat map presentation of Mixl1 responsive genes upregulated with Day 1 (early) and Day 3 (late) induction of Mixl1 activity in comparison to NoDox treatment. Gene Ontology enrichment analysis of the upregulated genes with Day 1 versus Day 3. E. Flow cytometry analysis of CXCR4+ cells in the differentiating EpiSCs with different timing of Mixl1 induction (N=3). The left panel is all CXCR4+ cells as a percentage of total cells. The right panel includes cells that were gated as highly positive for CXCR4. F. Expression of Sox17 transcript in MixL1WT, MixL1KO or MixL1Dox mouse EpiSCs with either No Dox treatment or MixL1 induction via Doxycycline at Day 0 (D0Dox), 1 (D1Dox), 2 (D2Dox) and 3 (D3Dox) of differentiation.

To generate EpiSC line in which *Mixl1* can be conditionally activated at different timing during differentiation, a FLAG-Mixl1 construct was integrated into the *Hprt* locus downstream of a tetracycline response element in the Mixl1KO ESCs. This ensured the built-in TET-On expression system to elicit time-controlled *Mixl1* expression in a Mixl1-LOF background (Figure 6A). For this study, we converted these ESCs to EpiSCs (Mixl1DOX) in vitro, in which induction by DOX enhanced the expression of FLAG-tagged *Mixl1* (Figure S8A, B). Mixl1DOX EpiSC expressed FLAG-*Mixl1* transcript in response to Doxycycline (Dox) treatment, and generated FLAG-MIXL1 protein (30KDa) in both monolayer culture and embryoid bodies (Figure S8B). However, attempts at detecting endogenous MIXL1 protein in the Mixl1WT cells were unsuccessful using two antibodies, which might be due to low levels of endogenous MIXL1 protein following transient Mixl1 expression or that expression is restricted to sub-set of cells responding to serum-induced differentiation. The findings from Mixl1DOX cells showed that the activation of *Mixl1* can be controlled in time which faithfully models the dynamic expression of *Mixl1* in the Mixl1WT EpiSCs.

We next characterized the impact of timed DOX-induced *Mixl1* expression on serum-induced differentiation of the Mixl1lDOX EpiSC line. DOX treatment commencing on different days (Day 0, 1, 2 and 3 = D0DOX, D1DOX, D2DOX and D3DOX respectively) led to the activation of FLAG-Mixl1 with expression levels peaking in the following 24-hour interval and expression then declined over the course of 120 h of differentiation (Figure 6B). Mixl1KO lines and untreated Mixl1DOX lines (NoDOX) did not display any activation of FLAG-Mixl1 (Figure S8A). While *Mixl1* transcripts were detected by qPCR in Mixl1KO and Mixl1DOX cell lines prior to differentiation (Figure 6D), these were the transcripts of the edited LOF alleles and therefore had no functional contribution.

A previous study of a library of EpiSC lines showed that early activation of Mixl1 expression (at Day 1-2 of differentiation) was associated with enhanced propensity of endoderm differentiation [17]. We compared the switch in response to Mixl1 activation using a transcriptomic dataset of bulk RNA-seq analysis of cells of the D1Dox, an early Mixl1 induction, and D3Dox, a late Mixl1 induction, at 24 h following induction. The Mixl1DOX cell line, differentiated for a similar time but without Doxycycline treatment (NoDox), was used as a control (Figure S8C). Principal component analysis of the 500 most variable genes displayed clear separation of the four conditions, except for two samples (Figure 6C). Although these samples could be excluded as outliers, it is possible that the variance is related to the technique used for generation of embryoid bodies. Therefore, these samples were included in all subsequent analyses. Over 90% of the variation, within the most variable 500 genes, is explained in the first two PC axes (Figure S8D, E). The negative PC1 axis separates the samples by Mixl1 induction as is exemplified by the high contribution of Mixl1 to the PC1 axis loadings. The orthogonal PC2 axis instead separated the samples by timepoints of differentiation.

Functional ontology analysis of the upregulated gene sets in response to early Mixl1 activation revealed enrichment of functionality of cell-substrate/matrix interaction, cell adhesion, tissue remodelling and epithelial morphogenesis whereas upregulated gene sets to late Mixl1 activation were enriched for ectoderm/neural differentiation (Figure 6D). The gene-sets with the enriched functional ontology that were upregulated in response to late *Mixl1* activation overlap with those that were downregulated in response to early *Mixl1* activation (Figure S8F), suggesting that MIXL1 may act to enhance the transcription of downstream genes for endoderm and mesendoderm differentiation and repress the gene activity for ectoderm differentiation. Further functional ontology analysis revealed that approximately a third of the upregulated early-Mixl1 response genes are shared with the late-Mixl1 response (Figure 6D, S8F). Gene ontology enrichment analysis of these common upregulated genes identified a combination of neurogenesis and cardiogenesis processes. Genes uniquely upregulated by early Mixl1 induction are enriched for cell adhesion and cell motility processes. Late induction of Mixl1, on the other hand, uniquely upregulated genes implicated in neural differentiation, a considerable shift from the early induction. Both early and late inductions reduced expression in genes for pattern specification and neural tube development, as well as those involved in BMP and WNT signalling pathways. Early induction reduced the expression of pattern specification process genes and neural cell type process genes, whereas late induction impacted on the expression of genes for epithelial cell fates. This difference in response gene sets aligned with previously observed outcome of differentiation, where late *Mixl1* activation resulted in a reduced proportion of endoderm as assessed by flow cytometry and higher expression of ectoderm markers as assessed by qPCR [38].

Flow cytometry was performed to quantify the proportion of CXCR4(CD184) positive definitive endoderm generated with different timing of *Mixl1* induction. The absence of *Mixl1* expression does not abrogate the emergence of cells expressing molecular markers of endoderm in vivo or in vitro [39]. Similarly, we observed a CXCR4+ population in NoDox culture (Figure 6E), the proportion of which varies between biological replicates of embryoid body differentiation. Activation of FLAG-Mixl1 resulted in an increased proportion of CXCR4+ cell proportions in line with the level of Sox17 expression (Figure 6F) suggesting that an elevated expression of endoderm-related genes is coupled with the enhanced generation of CXCR4+ endoderm. Furthermore, D1Dox EpiSC, which replicates the Mixl1-early phenotype of Mixl1WT EpiSCs, showed a greater enrichment of CXCR4+ endoderm particularly of a CXCR4-high population, compared to all other induction schedules (Figure 6E). A CXCR4-high population has not been previously reported during endoderm differentiation in vitro, and the implication or the identity of this CXCR4-high population is yet unexplored. Finally, we showed that the differential expression of endodermal gene (i.e., Sox17) within bulk populations among the EpiSC lines was reflecting a shift in the proportion of definitive endoderm cells in the Day 5 culture (Figure 6F). Taken together, these results point to a causal link between the timing of *Mixl1* expression and the enhanced propensity of endoderm differentiation.

## Discussion

In this study, we identified the putative mesendoderm progenitor cells (pMEP) in the gastrulating mouse embryo by computational analysis. Single cell RNA-sequencing of gastrulating embryos identified a unique gene set including the lineage-specific mesoderm and endoderm genes and genes that are expressed in the mesendoderm derived from the mouse and human pluripotent stem cells. By imputing the combined activity scores of this gene set, populations of pMEP are identified and localized to the epiblast and the primitive streak of the E7.0-E7.5 embryo. We next traced the MIXL1GFP positive cells in the Delmix mouse model to affirm the location of the pMEPs in mouse embryos at E7.0 - E7.75. This cell population is first localized in the proximal epiblast and the posterior region of the primitive streak and later in the distal epiblast and the anterior region of the primitive streak. Although the pattern of *MIXL1* expression in human embryos is yet undefined, *MIXL1* transcript has been identified in the posterior epiblast of the gastrulating embryo of the non-human primate cynomolgus monkey [40] where MIXL1 expression overlaps with that of *EOMES* and *BRACHYURY*. Single-cell and spatial transcriptomics studies of other non-human primates identify similar cell populations that are localised to the posterior region of the gastrulating embryo [41–43].

By tracing the lineage contribution of Mixl1GFP cells in the embryo, we showed that MixI1 marked cells can contribute to both mesoderm and endoderm. In addition, we identified descendants of Mesp1Cre-activated GFP+ cells in the gut endoderm of E8.5 mouse embryo. Subsets of the *Mixl1+/Mesp1+* cells may therefore constitute the putative mesendoderm progenitor. *Eomes*-expressing cells give rise to *Mixl1-* and *Mesp1-*positive cells that go on to populate the mesoderm and endoderm [44]. As *Mesp1* and *Mixl1* expressing cells co-exist in the primitive streak, it will be interesting to explore if this subset of cells that were Mesp1-active in the foregut are the same cells that display perduring Mixl1GFP expression in E7.75-E8.5 embryo. For such lineage tracing study, a combination of multiple lineage reporters in the same cells would be required.

The mechanism underpinning this mode of lineage allocation of the pMEPs may lie in the response to different levels of Nodal and BMP activity perceived by cells as they ingress at the primitive streak[45]. In addition, the relative activity of the effectors, pSMAD1/5 and pSMAD2/3, for BMP and Nodal respectively, may modulate the expression of their target genes, resulting in differential activity of lineage determinants of the mesoderm versus the endoderm. The BMP/Nodal gradient may also activate a different level of *T* and *Eomes* activity, leading to the specification of mesoderm lineage accompanied by high *T* activity and endoderm lineage by high *Eomes* activity [4, 46–49].

To guide the choice of lineage cell fate, timely expression of unique sets of transcription factors are induced in the primitive streak and adjacent epiblast. Among these is the Mix Paired-Like homeobox gene, *Mixl1,* which marks the primitive streak in the mouse [19, 50, 51]. Previous studies have identified a potential role of Mixl1 in guiding the lineage choices during germ layer differentiation [23, 52]. However, the role of Mixl1 in guiding lineage diversification of the pMEPs has not been elucidated. In this study, we showed that early activation of *Mixl1* promotes endoderm differentiation of the epiblast stem cells, whereas late or sustained activation of Mixl1 predisposes mesoderm and ectoderm differentiation. The impact of different timing of *Mixl1* activation on cell fate choices is reflected in the differential expression of lineage-related gene sets during the differentiation of the epiblast stem cells. These observations raise the possibility that Mixl1 is acting as the triage factor for lineage diversification of the putative mesendoderm progenitors. Early activation upon exit from pluripotency followed by expeditious downregulation of *Mixl1* would guide the progenitor cells to endoderm differentiation while belated activation and sustained *Mixl1* activity may channel the progenitors to the mesoderm lineage. Further study of other MEP-related genes may discover other genetic determinants of lineage choices that act in concert to mediate lineage diversification of the mesendoderm progenitors in vivo and in vitro.

## Materials and Methods

## RESOURCE AVAILABILITY

### Lead Contact

Further information and requests for resources and reagents should be directed to and will be fulfilled by the lead contact, PPLT.

### Data and Code Availability

- Single-cell RNA-seq data will be deposited at GEO and will be publicly available as of the date of publication. Microscopy data reported in this paper will be shared by the lead contact upon request.
- All code will be made publicly available as of the date of publication.
- Any additional information required to reanalyze the data reported in this paper is available from the lead contact upon request.

## EXPERIMENTAL MODEL AND SUBJECT DETAILS

### Mice

Mesp1Cre [31], ZEG [53], Delmix [23] and ARC strains of mice were used.

Animal experimentation was performed in compliance with animal ethics and welfare guidelines regulated by Children’s Medical Research Institute/ Children’s Hospital at Westmead Animal Ethics Committee, protocol number C364.

### Cell Lines

The HES3 Mixl1:GFP cell line (female) was cultured on hESC-qualified Matrigel (Corning) coated plates using mTESR plus media (StemCell Technologies) [34]. Cells were split twice weekly at 70% confluency using ReleSR (StemCell Technologies) for maintenance.

A2Lox.cre mES cell lines (male) were maintained on plates coated with 0.1% gelatin and non-commercial, irradiated E13.5 mouse embryonic fibroblast (MEF) feeders (at a density of 1.7×10^4^ cells/cm^2^) in ESC medium consisting of Dulbecco’s Modified Eagle’s Medium (DMEM; Gibco) supplemented with 12% heat-inactivated fetal calf serum (Gibco), 1× MEM Non-Essential Amino Acid Solution (NEAA; Gibco), 1× nucleosides, 1000 U Mouse Leukemia Inhibitory Factor Medium Supplement (LIF; ESGRO) and 0.1 mM 2-mercaptoethanol (Sigma). The composition of 100× nucleosides supplement is as follows: 0.8 g/L Adenosine (Sigma), 0.73 g/L Cytidine (Sigma), 0.85 g/L Guanosine (Sigma), 0.73 g/L Uridine (Sigma) and 0.24 g/L Thymidine (Sigma). Mouse ESC cultures were routinely passaged at 50-70% confluence using TrypLE Select (Gibco).

MixL1KO and MixL1Dox EpiSCs were maintained in EpiSC-medium consisting of KnockOut DMEM (Gibco) supplemented with 20% Knockout Serum Replacement (Gibco), 1% Non-essential amino-acid supplement, 1% Glutamax (Gibco), 0.1 μM 2-Mercaptoethanol, 10 ng/mL recombinant Human basic fibroblast growth factor (FGF) protein (Carrier Free) (R&D Systems) and 20 ng/mL recombinant human/mouse/rat Activin A protein (Carrier Free) (R&D Systems). Media was changed daily, and cells passaged at 70% confluency using 2 mg/mL Collagenase IV (Gibco) resuspended in KnockOut^TM^ DMEM to detach colonies followed by single cell dissociation with TrypLE Select (Gibco). EpiSCs were seeded at 5×10^4^ cells/cm^2^ on plates coated with 0.1% gelatin and seeded with MEF feeders as described for the A2LoxCre cell line above. For the first 24 hours, EpiSC-medium was supplemented with 10 μM Y-27632 ROCK inhibitor (TOCRIS). All cell cultures were maintained at 37 °C, 5% CO_2_ in air and 95% humidity. Cell lines routinely tested negative for mycoplasma infection.

### Single-cell RNA Sequencing Quantification and Statistical Analysis Read alignment and expression count table generation

From the sequencing results of the 10x Chromium experiments, the unique molecular identifiers, cell barcodes, and the genomic reads were extracted using Cell Ranger with default parameters (v3.1, 10x Genomics). The extracted reads were aligned against the annotated mouse genome, including the protein and non-coding transcripts (GRCh38, GENCODE v27)[54]. The reads with the same cell barcode and unique molecular identifier were collapsed to a unique transcript, generating the count matrix where columns correspond to single cells and rows correspond to transcripts. To remove potentially unhealthy or suboptimal cells, cell filtering was performed using the number of reads, the proportion of genes expressed, and the fraction of mitochondrial reads as criteria. Specifically, cells with greater than 99% of genes not expressed and 10% of mitochondrial gene expression were removed. Transcripts from mitochondrial- and ribosomal-protein coding genes were discarded for downstream analyses such as embedding and clustering, because they are typically known to be highly expressed irrespective of biological identity.

### Doublet detection and filtering

The presence of multiplets in single-cell data can arise from incomplete dissociation of single cells meaning that more than one cell can be encapsulated in GEMs. DoubletFinder, an algorithm to detect multiplets in single-cell data, was used to remove potential doublets or multiplets from each biological batch at a threshold of 10% [55].

### Dimensionality reduction, clustering, and cell-type annotation

To embed the single-cell transcriptomes into the latent space, we first performed principal component (PC) analysis on the binomial Pearson’s residuals calculated using the scry package [56] on the feature-selected count matrix. Feature selection was performed using scry, and the top 1500 genes were used. Using the first 40 PCs, the reduced embeddings were generated by using fftRtsne, the FFT-accelerated interpolation based tSNE from the FIt-SNE package [57]. The first 35 PCs were used to construct the shared nearest neighbor graph using the default arguments of the buildSNNGraph function in the scran R package [58] with which the graph-based Louvain clustering was performed to generate the final clusters. Cell identity analysis was performed using the Cepo R package [27, 58]. Finally, annotation of the clusters from the in-house datasets was performed by manual assignment of cell types through inspection of cell identity genes.

### Detection of cell identity genes and visualization of gene expression

Normalized gene expression was visualized as a dot plot or split violin and box plots. Within the box plots, the median is denoted as red circles. The box denotes the inter-quartile range. Outliers have been omitted to facilitate visualization. The student’s *t*-test was used to compare the difference in mean gene expression between healthy and diseased cells. ns: p-value > 0.05; *: p-value <= 0.05; **: p-value <= 0.01; ***: p-value <= 0.001; ***: p-value <= 0.0001. The ggplot2 and introdataviz R packages were used for visualization [59, 60].

### Computational approach to predict population of biprogenitors

The mesendodermal biprogenitors were predicted using k-nearest neighbours (k-NN) algorithm, which is a supervised machine learning algorithm predominantly used for classification [61]. We first individually trained two KNN classifiers to predict either the mesodermal or the endodermal lineage cells. For each classifier, the top 200 cell identity genes from the Cepo algorithm for the mesoderm and endoderm populations were used. We initialised the classifier and returned class probabilities for each cell. Because we hypothesize that biprogenitors are a rare subclass of population, we created an ensemble framework that would broaden our search of cells that can be identified as biprogenitors by varying the number of k-neighbours (5 to 20) and probabilities (0.6 to 0.8). We then performed a search of mesodermal or endodermal cells, training on mesodermal cells with predicted class probabilities between 0.6 and 0.8. Cells with high class probabilities (>0.8) were excluded to prioritise predictions of cells that have a mixed profile of mesendodermal characteristics. Finally, we assigned cells that have been classified as both mesoderm and endoderm cells as mesendodermal biprogenitors across all combinations of the permutations.

### Identification of markers of biprogenitor cells

To accurately identify markers of biprogenitors, we performed an independent marker analysis of biprogenitors using our in-house dataset and from a previous study that profiled mouse gastrulation at single-cell resolution [62]. The mesendodermal progenitors were predicted following the same framework described in “Computational approach to predict population of biprogenitors” on the two datasets. The top 500 cell identity genes of the biprogenitor populations were identified using Cepo [27]. We further defined a set of genes that are highly correlated with T/ Brachyury expression at E7.0 using the iTranscriptome database [63]. Finally, we defined the intersection of these three groups as the high-confidence markers of MEPs, which led to a total of 20 genes. The MEP signature expression was generated by calculating the average gene expression in log2(FPKM) from the iTranscriptome database [12, 63] of the MEP marker genes (either from the Cepo-derived set or the refined 20 signature MEP genes) and visualised as cornplots using the ggplot2 package [60].

### Spatial allocation of single cells to the spatial mouse embryo

To map the spatial locations of the predicted biprogenitors to the mouse embryo, we used our scRNA-seq data from the mouse embryos at E7.0 and the corresponding spatial transcriptomes of mouse embryos at the E7.0 developmental stage generated using the GEO-seq protocol [64]. We used the union of the top 75 highly variable genes from each of the spatial domains to map single cells to their spatial locations. We then assigned each single cell to the best matched spatial location based on Spearman’s correlation of the expression of the selected genes. In particular, we calculated the purity of spatial locations for the MEPs based on the assigned single cells. Specifically, we define the purity of a spatial location 𝑙(𝑙 = 1, . . ., 𝐿) as 𝑝*urit*𝑦_𝑙_ = 𝑚*ax*(*𝑝*_1_, *𝑝*_2_, . . ., *𝑃_M_*), where 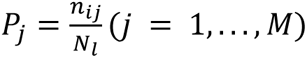 is the percentage of cell type 𝑗 at that spatial location of the embryo, *𝑛_𝑙𝑗_* is the number of cells in that cell type assigned to 𝑙 and 𝑁_𝑖_ is the total number of cells assigned to 𝑙𝑙.

### Mouse Embryos

#### Mouse embryo collection

Mixl1+/GFP mice were crossed with ARC mice to collect embryos at E7.0, E7.5 days post coitum. Mesp1-Cre mice were crossed with ZEG reporter mice [53] to collect the Mesp1Cre::ZEG embryos at E7, E7.5, E8, E8.5, E9.5 days post coitum. Imaging of the collected mouse embryos to screen for positive GFP was performed on Zeiss Lumar microscope.

#### Immunofluorescence of mouse embryos

Mixl1^+/GFP^ mouse embryos were collected at E7.5 days post coitum [23]. Whole mount fluorescent immunostaining of mouse embryos was performed as described in Masamsetti et.al. 2022 [65]. Briefly, collected embryos were fixed in 4% paraformaldehyde (PFA) (Scharlau) for at least 1 hour. Embryos were then washed in Dulbecco’s phosphate buffered saline (PBS) (Sigma) before permeabilizing with 0.4% Triton X-100 (Sigma). Embryos were then blocked in 1% BSA (Sigma), PBS with 0.1% Triton X-100 (PBST) for 1 hour, followed by incubation with primary antibody, FOXA2 (Abcam) 1:200 and SOX17 (R&D Systems) 1:200, overnight. Embryos were washed thrice in PBST and then incubated with respective secondary antibodies for 2 hours. These embryos were then washed thrice with PBST to remove unbound secondary antibodies and then cleared using FUNGI solution [66].

### hESC Micropatterning

#### Micropattern preparation

Micropattern coverslips were fabricated based on the protocol of Lee et. al., 2016, with some modifications. In brief, glass coverslips were sonicated in 70% ethanol and in deionized water. The clean coverslips were sequentially incubated in 0.5% (3-aminopropyl)triethoxysilane (APTS) (Sigma) for 3 min, 0.5% glutaraldehyde (Sigma) for 30 min. After air drying, the coverslip was deposited on a 20µL drop made of 10% acrylamide (Sigma), 0.87% N-N’-Methylenebisacrylamide (Sigma), 0.1% ammonium persulfate (Sigma) and 0.1% N,N,N′,N′-Tetramethylethylenediamine (Sigma), to make the gel at a stiffness of 100KPa. After the stiffness droplet was semi-solidified, the whole system was submerged into 70% ethanol, resulting in a smooth polyacrylamide gel forming. Gelled coverslips were sequentially coated with hydrazine hydrate (64% hydrazine; Fisher Scientific) for 1 hour and 5% glacial acetic acid (Ajax Finechem) for 1 hour.

To generate polydimethylsiloxane (PDMS) stamp, SYLGARD™ 184 Silicone Elastomer Curing Agent and Base (Dow) were mixed at a 1:10 ratio before loading to the stamp mould, provided by the Kilian Lab at the University of New South Wales. Next, the solidified PDMS stamp was coated with 25 μg/mL vitronectin (Life Technologies) and 3.5 mg/mL sodium periodate (Sigma) for 30 min. After air-drying the stamp, patterned vitronectin was stamped onto the gelled coverslip at 0.343N for 1 min. Stamped gels were stored overnight in PBS + 1% Penicillin-Streptomycin (Sigma) and 0.25 µg/mL Amphotericin B solution (Sigma) at 4 °C.

#### Mesendoderm differentiation of human ESCs on Micropatterns

Differentiation of the cells on micropatterned coverslips was a protocol adapted from Warmflash et al. 2014 [67]. HES3 Mixl1:GFP cells were passaged as single cells with StemPro Accutase Cell Dissociation Reagent (ThermoFisher Scientific) and seeded on micropatterned coverslips at a density of 2.5x10^5^ cells/cm^2^ with 10μM Y-27632 dihydrochloride Rock inhibitor (TOCRIS) in mTeSR Plus (StemCell Technologies) into a total volume of 0.5 mL per well of a 24-well plate. Cells were incubated at 37 °C with 5% CO_2_. After 24 hours, micropatterns were incubated with either 50 ng/mL recombinant human BMP-4 (R&D Systems) or 3 µM CHIR99021 (StemCell Technologies) and 100 ng/ml recombinant h/m/rActivin A (R&D Systems) in mTeSR Plus and incubated for a further 48 hours at 37 °C with 5% CO_2_.

#### Immunostaining of Micropatterns

The cells grown on micropatterns were washed with PBS, fixed in 4% paraformaldehyde in PBS at RT for 20 min. They were washed with PBS twice and then permeabilised with 0.1% Triton X-100 (Sigma) in PBS (PBST) at RT for 1 hour. The cells were blocked with 3% bovine serum albumin (Sigma Aldrich) in PBST at room temperature for 1 hour. They were incubated with primary antibody in PBST at 4 °C overnight. Primary antibodies includeFOXA2 at 1:300 (Abcam), T/Brachyury at 1:20 (R&D Systems) and SOX17 at 1:20 (R&D Systems. On the next day, the cells were washed with PBST thrice, then incubated with corresponding secondary antibodies at 1:700 in PBST RT for 1 hour. The cell nuclei were stained with DAPI (1 μg/ml) (ThermoFisher Scientific) in PBS for 10 min at RT, and then washed twice more with PBS. Cells were mounted with Fluoromount-G (ThermoFisher Scientific).

### Microscopy Imaging and Analysis

#### Confocal microscopy

Immunostained Micropatterned coverslips and whole mount mice were imaged on Zeiss AiryScan LSM 880 confocal microscope. Zen and Imaris 3D rendering softwares were used for visualization and 3D representation of images.

#### Confocal imaging of mouse embryos

Confocal Z-stacks were acquired every 8µm through the depth of the embryo and collapsed to obtain maximum intensity projections. Single cell co-expression with multiple protein expression was observed and identified on individual optical sections. Imaris software was used for 3D visualization and analysis of confocal stacks. Optical sections of the 3D embryo were recorded using ortho/oblique functions in Imaris software.

#### Confocal imaging of micropatterns

Confocal Z-stacks were acquired every 8µm through the depth of the micropattern and collapsed to obtain maximum intensity projections. Single cell co-expression with multiple protein expression was observed and identified on individual optical sections.

#### Micropattern image analysis

The nuclei from micropatterned images taken were segmented using the StarDist method [68] via QuPath v0.4.3 [69] using default parameters and the *dsb2018_heavy_augment.pb* model. For the quantification of positively stained nuclei, QuPath was used, where the algorithm for positive detection was based on a pixel classifier trained on representative images with specific annotations.

### Epiblast Stem Cells

#### Generation of Mixl1 KO and Mixl1 DOX EpiSC lines

A2Lox.Cre mouse ESCs [70] was used to generate Mixl1 KO. Briefly, CRISPR/Cas9 was used to edit the *Mixl1* sequence at the first exon. Two guide RNAs (gRNA) targeting the first exon of *Mixl1* were cloned into an all-in-one plasmid pSpCas9(BB)-2A-GFP (pX458) (Addgene Plasmid #48138) [71]. 5 × 10^6^ mouse ESCs were transfected with 5 µg plasmid in a 100 µL volume using Neon Transfection System (ThermoFisher Scientific) with the following pulse set up: Voltage 1200 V, Width = 2 ms, Number = 2. Cells were seeded in three 10 cm dishes precoated with 0.1% gelatin and mouse embryonic fibroblast (MEF) feeders (1.7 × 10^4^ cells/cm^2^) for recovery. GFP expression was monitored daily until 96 h when colonies were large enough to be manually picked. 18 clones per gRNA were picked, transferred into a well of 96-well plate containing 40 µL of TrypLE Select (Gibco), incubated for 5 min at room temperature and finally, the single-cell suspension was transferred into a well of 96-well plate containing 150 µL of ESC media and pre-seeded with MEFs. Individual clones were split 3 times on MEFs then feeder-free, on 0.1% gelatin-coated plates, for sequencing. Cells (grown on MEFs) were frozen during the sequencing analysis.

Clones were sequenced for each gRNA. Regions flanking the predicted edited region were amplified by PCR and run on a 1% agarose gel. Clones displaying one band, or three bands, were discarded. Clones with two bands had whole DNA extract submitted to the first round of sequencing. Sequences were analysed using Tracking of insertions and deletions (INDELs) by decomposition (TIDE) online tool (http://shinyapps.datacurators.nl/tide/) [72]. Clones displaying a biallelic frameshift mutation were submitted to a second round of sequencing, where the PCR fragment was cloned into pGEM^®^-T Easy plasmid (Promega). White colonies were selected and further grown to amplify the DNA to be sequenced. A total of 8 plasmids per clone were sent for sequencing. Clones with a 50/50 ratio of both alleles retrieved and successfully sequenced were selected for our study.

E7.5 R129 mouse RNA extract was reverse transcribed using SuperScript III Reverse Transcriptase (ThermoFisher Scientific) as per manufacturer protocol to produce cDNA. *Mixl1* cDNA was amplified from E7.5 R129 mouse cDNA using the following primers: 5’-GGGGACAAGTTTGTACAAAAAAGCAGGCTTCGCCGCAGCAGGGTCCCAG-3’ and 5’-GGGGACCACTTTGTACAAGAAAGCTGGGTCTCAGAAGTTACCTAAGGC-3’. The amplified sequence was then run on a 1% agarose gel, column purified with Wizard SV Gel and PCR Clean-Up System (Promega) and cloned into an entry vector. To generate the p2Lox-FlagMixl1 plasmid (Gateway strategy – Invitrogen), the entry vector was recombined with a donor vector, containing the FLAG tag and the ATG sequence to rescue neomycin resistance in the A2Lox.Cre mESC line [70]. The p2Lox-FlagMixl1 plasmid was electroporated using the previously described strategy. Cells were selected using Neomycin for 2 weeks after which clones were picked for genetic screening and sequencing.

#### Epiblast stem cell derivation from mouse embryonic stem cells

Chimera and mouse EpiSC generation from Mixl1 KO and Mixl1 DOX ESC were performed as described previously[37] . Briefly, ESCs grown for 48 h in 10% FCS + 2i/LIF were injected into 8-cell mouse embryos, transferred into pseudopregnant mice uterine horns and allowed to develop. Epiblasts were collected from E7.5 embryos and seeded on mouse embryonic fibroblasts (MEFs) in EpiSC medium. Once EpiSCs were stably growing, Geneticin (Gibco) was added to the medium for 4 passages to select against host embryo cells.

#### Serum-induced differentiation of mouse epiblast stem cells in embryoid bodies

Embryoid bodies (EB) were maintained in EB-medium consisting of Dulbecco’s Modified Eagle’s Medium (Gibco) supplemented with 15% Fetal Calf Serum (Gibco), 1% Non-essential amino-acid supplement (Gibco) and 0.1 μM 2-Mercaptoethanol (Sigma). EpiSC colonies were collected after Collagenase IV (Gibco) treatment and transferred to EB-medium within non-tissue culture treated plates and placed at 37 °C, 5% CO_2_ on an orbital-shaker under slight rotation to prevent single EBs from settling to the bottom of the plate or attaching to each other. Media was replenished every 48 h, unless otherwise noted. Doxycycline induction, where appropriate, was achieved by adding 1 μg/mL Doxycycline (Sigma) to the culture.

#### In vitro conversion of mouse embryonic stem cells to Epiblast stem cells

Mouse embryonic stem cells were converted to Epiblast-Like stem cells (EpiLC) by plating on fibronectin and cultured in Ndiff media supplemented with Activin (20ng/ml) and FGF (12ng/ml) treatment for 48hours. These EpiLCs were then converted to Epiblast stem cells by culturing in EpiSc media {Knock out serum (20%), NEAA (1%), Glutamax (1%), β−Mercaptoethanol added to Knock Out DMEM)} supplemented with Activin (20ng/ml) +FGF (10ng/ml). Converted Epiblast stem cells (Conv. EpiSCs) were passaged for at least 9 passages, before performing experiments.

#### Embryoid body generation from converted Epiblast stem cells (Conv. EpiSCs)

Conv. EpiSCs were dissociated from plates by collagenase treatment. The colonies were grown in suspension culture on a 10cm dish using embryoid body making media {Fetal calf serum (15%), NEAA (1%), Glutamax (1%), β−Mercaptoethanol added to Knock Out DMEM}. Embryoid bodies were cultured in suspension for differentiation. On Day 3, embryoid bodies were collected, washed with PBS, fixed with 4%PFA and immunostained with respective antibodies (eg: Foxa2, SOX17).

#### Flow cytometry and Fluorescence-activated cell sorting (FACS)

Embryoid bodies for quantification of endoderm propensity were dissociated with TryPLE Select (ThermoFisher Scientific). CXCR4 (CD184) staining for endoderm propensity quantification was performed using phycoerythrin (PE)-conjugated rat anti-mouse CXCR4 antibody (R&D Systems) in Dulbecco’s phosphate-buffered saline with 10% Fetal Calf Serum (Flow cytometry buffer) for 45 minutes at room temperature and washed thrice in Flow cytometry buffer. Rat IgG2B Phycoerythrin isotype control (R&D) was used as isotype control. Cells were sorted by FACS Canto, quantified using the FACS Diva software (BD). Data figures were created using FlowJo Software (BD).

### RNA extraction

Total RNA from flash frozen samples was extracted using ISOLATE II RNA Mini Kit as per manufacturer recommendations (Bioline). In short, samples were thawed on ice, lysed and passed through a column purification process and DNase treatment. Precipitated RNA was eluted into RNase-free water. RNA concentration, DNA and ethanol contamination were measured using the NanoDrop (Thermo Fisher) and adjusted using RNase-free water as required.

### RNA-seq library preparation and sequencing

RNA-seq libraries were prepared from extracted RNA by Genewiz (Azenta, China) using the NEBNext Ultra II RNA Library Preparation Kit (NEB, E7770) as per manufacturer protocol and 150 bp paired-end sequenced on the HiSeq 4000 (Illumina).

### RNA-seq

150 bp paired-end reads were trimmed using Trim Galore [73] in paired mode ‘--paired’ keeping reads with at least 50 bp ‘--length 50’. Trimmed reads were aligned to the GRCm38/mm10 mouse reference genome using STAR aligner [74] with the parameter ‘--outSAMtype BAM SortedByCoordinat’ to output a coordinate-sorted BAM file for downstream processing. Transcript abundances were estimated from genome aligned sequences using featureCounts [75] in paired-end mode keeping read pairs in which both reads aligned but excluding chimeric assignments ‘-p -B’. Differential expression and statistical analysis were done using the DESeq2 library [76]. Differentially expressed genes were identified based on the following criteria: absolute Fold change >= 2 and adjusted *P*-value < 0.05. Biological processes enriched in differentially expressed RNA-seq gene sets were identified using the clusterProfiler library [77].

## QUANTIFICATION AND STATSITICAL ANALYSIS

Statistical analyses were performed using Graph Pad Prism 9. For comparison between two mean values a 2-tailed Student’s t-test was used to calculate statistical significance. Quantitative data are shown as mean +/- standard error and are considered significant when p<0.05 (*p <0.05, **p<0.01, *p<0.001).

## Acknowledgements

This work was supported by Australian Research Council (Discover Project grant DP 190102793, PPLT), National Health and Medical Research Council (Senior Principal Research Fellowship App1110751 PPLT) and the late Mr James Fairfax, Bridgestar Pty Ltd. We are grateful for Ashleigh Hope and Grady Smith for their technical assistance. We thank Blood Development Laboratory, Murdoch Children’s Research Institute for providing the Human: HES3 Mixl1: GFP.

## Author Contributions

VPM, NSN, NS, MW, HK, JS, JBS, PO conducted the experiments. VPM and PT designed and supervised the experiments. HJK, NA performed bioinformatic analysis. PY supervised the bioinformatic analysis. VPM and PT wrote the paper. All authors edited the paper. PT led the study.

